# Long-read Data Revealed Structural Diversity in Human Centromere Sequences

**DOI:** 10.1101/784785

**Authors:** Yuta Suzuki, Gene Myers, Shinichi Morishita

## Abstract

Centromeres invariably serve as the loci of kinetochore assembly in all eukaryotic cells, but their underlying DNA sequences evolve rapidly. Human centromeres are characterized by their extremely repetitive structures, *i.e.*, higher-order repeats, rendering the region one of the most difficult parts of the genome to assess. Consequently, our understanding of centromere sequence variations across human populations is limited. Here, we analyzed chromosomes 11, 17, and X using long sequencing reads of two European and two Asian genomes, and our results show that human centromere sequences exhibit substantial structural diversity, harboring many novel variant higher-order repeats specific to individuals, while frequent single-nucleotide variants are largely conserved. Our findings add another dimension to our knowledge of centromeres, challenging the notion of stable human centromeres. The discovery of such diversity prompts further deep sequencing of human populations to understand the true nature of sequence evolution in human centromeres.

## Main

Centromeres have been one of the most mysterious parts of the human genome since they were characterized, in the 1970s, as large tracts of 171 bp strings called alpha-satellite monomers^1, 2^. With a growing body of evidence suggesting their relevance to human diseases as sources of genomic instability or as repositories of haplotypes containing causative mutations^3–8^, it has become more important to investigate the underlying sequence variations in centromeres^9, 10^.

Human centromeres have nested repeat structures. Namely, a series of distinctively divergent alpha-satellite monomers compose a larger unit called higher-order repeat (HOR) unit, and copies of an HOR unit are tandemly arranged thousands of times to form large, homogeneous HOR arrays. While HOR units are chromosome-specific and consist of two to 34 alpha-satellite monomers, copies of an HOR unit are almost identical (95 ~ 100%) within a chromosome (Figure 1a)^11–17^. The total HOR array length in each chromosome is known to differ dramatically among individuals^7, 18^ or across human populations^19, 20^. In addition to array length, other types of variation are known to exist within an HOR array, such as structurally variant HORs that consist of different numbers and/or types of alpha-satellite monomers^20–23^ and also single-nucleotide variations (SNVs) between the HOR units^20, 24, 25^. One of the major driving forces leading to such centromeric variation has been thought to be structural alterations such as unequal crossovers and/or gene conversions^26, 27^.

**Figure 1.**
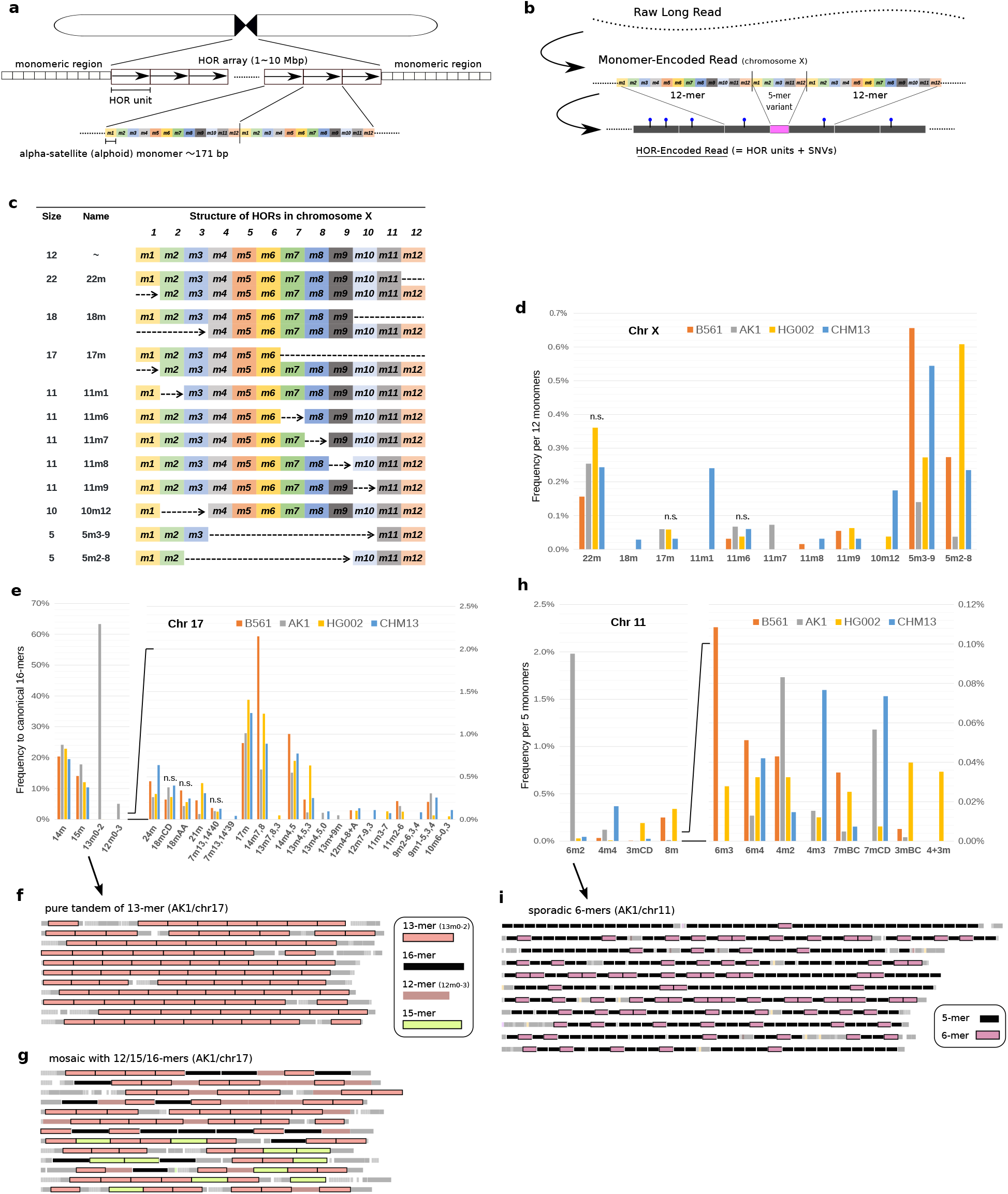
Comprehensive detection of variant HORs in centromeric arrays. (a) Schematics of a typical DNA sequence structure of human centromeres. The entire region consists mostly of alphoid monomers of 171 bp long. The core centromeric regions (several Mb) with HOR structure are sandwiched by the pericentromeric (monomeric) regions, where monomers are arranged tandemly without HOR. (b) Steps for HOR-encoding of long reads. Monomer-encoded reads were obtained by aligning monomer sequences into raw long reads, then frequent patterns of assigned monomers were considered HORs. The blue pins indicate the mismatches recorded in HOR-encoded reads, which contain both SNVs and sequencing errors. (c) Structures of canonical and variant HORs detected in chromosome X. The rectangles represent the presence of corresponding alphoid monomers. No gap is allowed between two constituent alphoid monomers to be detected as HORs. (d, e, h) Relative frequencies of detected variant HORs for four samples, B561 (orange), AK1 (gray), HG002 (yellow), and CHM13 (blue), in (d) chromosomes X, (e) 11, and (h) 17. “n.s.” (not significant) indicates the variants whose frequency was not significantly different across the samples. (f, g) The 24 longest reads containing the 13-mer variant HORs (red rectangles) from AK1 showed purely tandem structures as in (f), or mosaicism with other variant HORs (brown 12-mers, light green 15-mers) and/or canonical 16-mer HORs (black) as in (g). Detected HORs are represented as solid rectangles and other unexplained monomers due to sequencing errors are in gray boxes, placed proportionally to their actual positions within reads. (i) The 11 longest reads containing the 6-mer variant 6m2 (pink rectangles) from AK1. While the variant seemed enriched in reads, their distribution was sporadic, at most three variants were found in tandem.

Previous studies have investigated centromeric sequence variations via traditional approaches such as restriction enzymes sensitive to alpha-satellite monomers, Southern blotting, or analysis of k-mers unique to centromeres in short reads obtained in the 1000 Genomes Project^28^, but their observations have remained indirect and were confined to specific types of variations due to technological limitations.

Recently, the advent of long-read sequencing technologies has paved the way for direct, comprehensive observation of sequence variations among various human populations^29–33^. While long-read sequencing was capable of yielding contiguous reference sequences of centromeres for several species^34, 35^, reconstruction of whole centromeric sequences for human genomes is still challenging due to their idiosyncratic repeat structures. To date, it has been achieved only for the X and Y chromosomes, which both exist in a haploid state^36, 37^. While reference-quality *de novo* assembly of such repetitive regions remains a demanding task involving substantial manual curation^37–39^, the use of unassembled long reads has promise for investigating centromere variations in a cost-effective manner^40^.

Therefore, we exploited a strategy of HOR encoding of unassembled, uncorrected long reads for comprehensive detection and quantification of variant HORs. The use of unassembled reads enabled us to analyze diploid samples. In addition, the uncorrected reads could address SNVs in the HORs in an unbiased way. Here, we revealed a hidden diversity of centromeric arrays in terms of variant HORs through analysis of long reads from four human samples of diverse origins. We identified many novel variant HORs including some specific to one or two samples, and even when variants were shared across the four, their observed frequencies were substantially different.

## Results

### Direct determination and quantification of HOR variants via HOR encoding of long reads

To investigate interindividual variations within the centromeric array, we analyzed single-molecule real-time sequencing reads from four samples: B561 (Japanese), AK1 (Korean)^30^, HG002 (Ashkenazi)^33^, and CHM13 (European)^31^. First, the long reads were preprocessed in silico to filter out the noncentromeric fraction, and the remaining reads were interpreted as a series of alphoid monomers using a catalogue of 1197 monomers, *i.e.*, represented as monomer-encoded reads (Figure 1b). Then the monomer-encoded reads were clustered based on the composition of the different types of monomers, and for each cluster of reads associated with one of the HOR arrays, we constructed a catalog of variant HORs by detecting frequent patterns in the monomer-encoded reads. Finally, HOR-encoded reads were obtained by automatically replacing these patterns with symbols representing HORs. Below, we focused on HOR arrays in chromosomes 11 (D11Z1), 17 (D17Z1), and X (DXZ1), which evolved from the 5-mer archetypal HOR, in which we expected more complex variations could be found than in the other chromosomes associated with the dimeric archetypes^16^. We avoided chromosomes 1, 5, 13, 14, 19, 21, and 22, where chromosome identity is obscured because of shared HOR patterns. For CHM13, we performed the same analysis using Oxford Nanopore Technologies (ONT) data^37^ for comparison with the PacBio data.

#### Diversity of HOR variant frequencies in four different samples

The detected variant HORs showed a wide diversity in terms of presence and abundance among the four samples.

In chromosome X, the canonical HOR consists of 12 monomers, and it was the most frequent pattern found in reads across the four datasets (98.0% ~ 99.1% of all HOR types). Other than the canonical HOR, 11 variant HORs whose sizes ranged from 5-mer to 22-mer were defined (Figure 1c, d, Supplementary Figure 1). While five variants were shared among the four samples, six were found in <4 samples. For example, the 18-mer, an 11-mer were specific to CHM13 and another 11-mer was found only in AK1. To understand which variant HORs exhibited notable variability even in the presence of a small number of false positives and false negatives in the detection of variant HORs, we performed chi-square tests against the models where the relative frequency of variants should be constant across four samples. At least eight variants showed significant variation (*p* < 0.01 after correction for the number of tests) among the samples (Supplementary Table 1).

Similarly, for chromosome 17, 21 distinct variants of sizes from 7-mer to 24-mer were detected (Figure 1e). Twelve of them were specific to <4 samples. Notably, the 13-mer (in which three monomers, the 10th, 11th, and 12th, were deleted from the 16-mer) was present at high frequency only in AK1 and was essentially missing in the other three samples. Thus, AK1 turned out to have haplotype II, which is known to be shared among 35% of individuals^25, 41^. Consequently, while all datasets agreed that the most frequent pattern was the HOR made up of 16 monomers, its densities were different, measuring 59.2%, 35.4%, 59.7%, and 57.1% (per 16 monomers), respectively, in B561, AK1, HG002, and CHM13(PacBio). Again, the distribution of variant HORs across the individual samples was far from uniform (Supplementary Figure 4, Supplementary Table 1).

In chromosome 11, the 5-mer canonical HOR^16^ was the most frequent species, representing 97.4% ~ 99.4% of all HOR types. In addition, 18 variant HORs were detected, 12 of which comprised the same repertoire of 5 monomers as the canonical HORs, while the other six variants exhibited slightly different compositions in terms of the monomers they contained (Supplementary Figure 3). As with the other chromosomes we investigated, these variant HORs were observed with significantly variable frequencies across the four samples (Figure 1h). The most prominent difference was observed for a 6-mer variant (6m2; deletion of the second monomer), which accounted for 1.98% of the HORs detected in AK1. The same 6-mer variant was present at very low frequencies in two European samples HG002 and CHM13, but it was completely missing in the Japanese sample, B561.

#### The different modes of local expansion of variant HORs

Because the variant HORs were detected in long reads, we could investigate the contexts within which they were found (Figure 1f, g, i). In general, we found several examples in which the same type of variant HOR was enriched in a read, which suggested that these variants might have expanded locally through a series of duplication events, rather than occurring independently. Moreover, different modes of expansion were in evidence depending on the type of variant HOR. For example, the characteristic 13-mer variant (13m0-2) detected in chromosome 17 of AK1 was found in tandem or interleaved with other HORs (Figure 1f, g). In contrast, the 6-mer variant (6m2) detected in chromosome 11 of AK1 was found in a sporadic manner (Figure 1i). Therefore, unlike the variant 13m0-2, 6m2 seemed incapable of independent tandem expansion, and some preference (*e.g.*, length) with respect to the unit of expansion may exist.

### Local evolutionary patterns within centromeric array by rare and specific variant HOR motifs

We searched the HOR-encoded long reads for rare variant HOR motifs that were conserved among samples. We excluded the possibility that such a rare motif would have emerged independently multiple times in centromeres, which is an implausible hypothesis according to the maximum-parsimony criterion. In addition, we observed at most two patterns in the context of such rare motifs within the long reads from each of the three diploid individuals, and a single pattern for CHM13, implying that these regions with rare motifs were homologous. Therefore, we could compare the regions around each motif to understand local sequence evolution patterns in centromeres.

For example, we observed a combination of 12-mer and 5-mer variant HORs in chromosome 11 and the reads with the motif could be aligned and clustered into a single group (homozygous) or two groups (heterozygous) in each sample, revealing at least six alternative loci across four samples (Figure 2a, Supplementary Figures 5, 6).

**Figure 2.**
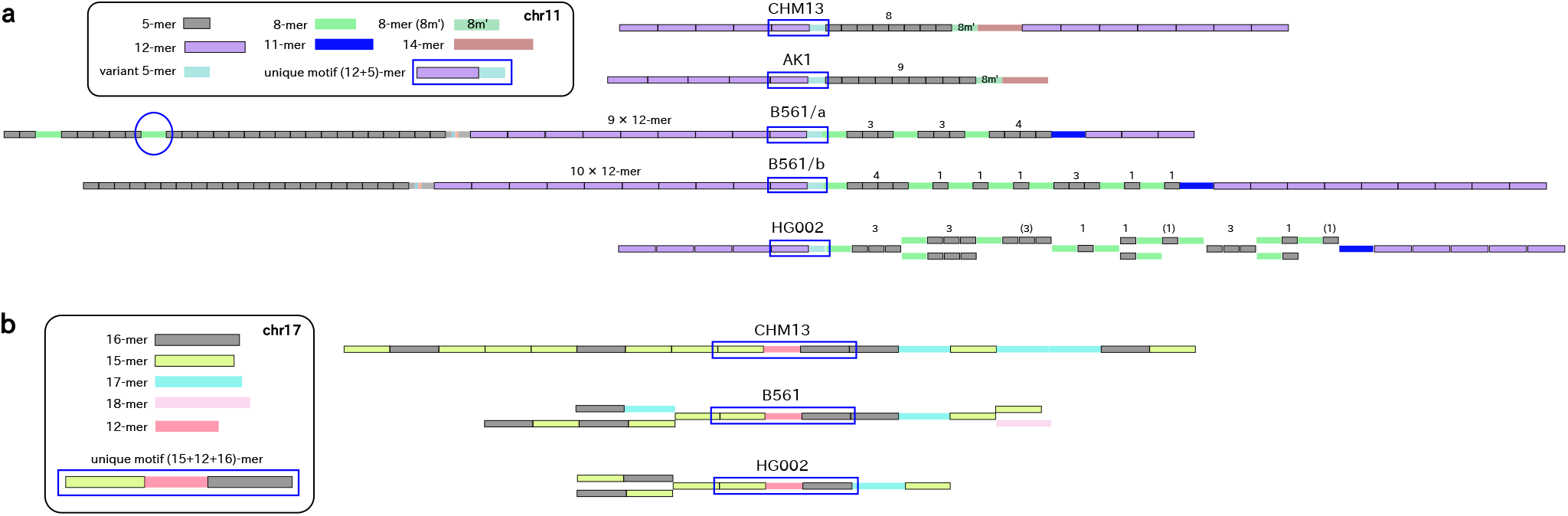
Patterns found around rare motifs of variant HORs. Canonical HORs and monomers are shown in grey, and each variant HOR was colored differently. (a) The specific combination of 12-mer and variant 5-mer (enclosed by blue lines) was found in all samples in similar but slightly different contexts. In CHM13 and AK1, there were another 8-mer plus 14-mer motif in downstream, but the number of canonical 5-mer HOR between them was differed by one unit. In B561, the differences found in upstream and downstream of the motif could be phased into two alternative patterns, suggesting they were two alternative alleles. In HG002, the alternative patterns in downstream could not be phased due to shorter read length, but explained by two loci. (b) The specific combination of 15-mer variant, 12-mer variant, and 16-mer canonical HOR in this order in three samples. While there was a single pattern in CHM13, two alternative patterns were found in upstream and/or downstream in B561 and HG002.

Although each cluster may not represent a single locus, we were able to compare homologous regions and identify structural alteration events around them. In B561, the downstream motifs were characterized by the presence of interleaving 8-mer variants, the 11-mer, and then, tandem 12-mers. The long reads suggested that the motif was located in two alternative loci with similar but clearly distinct structures (Figure 2a). In HG002, similar patterns were observed and at least two alternative loci were suggested, but we could not phase them due to insufficient read length. Reasonably, there were structural alteration events involving the variant HORs, and both B561 and HG002 were heterozygous in terms of HOR-level variation. In AK1 and CHM13, which were depleted of the 8-mer variant, the downstream motifs were tandem canonical 5-mers (8 copies in CHM13, 9 copies in AK1), followed by the combination of another type of 8-mer variant and the 14-mer variant. These differential patterns also suggested that the expansion or contraction had occurred in one of these individuals.

Similarly, we identified an additional motif comprising a 15-mer, 12-mer, and 16-mer in chromosome 17 (Figure 2b, Supplementary Figure 7). While all the reads with this motif suggested a single locus in haploid CHM13, two loci were found for each of the two diploid samples (B561 and HG002), and the pattern was missing altogether in AK1. Thus, in total, the motif appeared in five alternative alleles, assuming that they were all homologous. Overall, these examples, as well as the previous examples, demonstrated the highly variable nature of the centromere sequences.

### How useful are HOR variants in assembling centromeric sequencing reads?

Given that centromeric arrays contain ample HOR variants, one might also expect that they would serve as unique anchors useful in genome assembly. Put differently, with enough HOR variants, centromeric arrays would no longer appear repetitive. This intuition needs to be examined carefully, because the vast majority of the HORs found in centromeric arrays are canonical, and HOR variants remain rare. For example, the fraction of variant HORs among all the detected HOR units in chromosome X was 0.9% ~ 2.0% across our four samples. Considering this low rate of variant HORs that serve as positional markers, we calculated the probability that a window of a given length was locatable, *i.e.*, contained multiple HOR variants that could determine the relative position of the region compared to others. We assumed that HOR variants occurred in centromeres according to the Bernoulli distribution. Then, in an array with 99% canonical 12-mers (mimicking the situation in chromosome X), the probabilities that the regions would be locatable were 2.4%, 8.3%, and 16.6%, respectively, for 50 kb, 100 kb, and 150 kb window sizes, leaving most regions unlocatable. In an array with 99% canonical 5-mers (similar to chromosome 11 in our data), the probabilities were 11.5%, 32.3%, 52.3%, respectively, for 50 kb, 100 kb, and 150 kb window sizes. Therefore, we speculate that much longer reads would be needed to assemble centromeric regions based solely on the presence of HOR variants.

### SNV landscapes on the canonical HORs

Next, we analyzed SNV distribution over the canonical HORs, i.e., 5-mers in chromosome 11, 16-mers in chromosome 17, and 12-mers in chromosome X. Here, we did not analyse indels because they cannot be called confidently within long reads. Although most of the alternative bases were observed with a certain frequency (approximately a few percent) due to remaining substitution errors in the long reads, we could spot prevalent SNV sites as prominent peaks in the plots (Figure 3a-d, Supplementary Figures 8, 9). For example, the number of SNVs detected with a frequency above 5% among the canonical 5-mers in chromosome 11 were 25, 17, 17, and 20, respectively, for the B561, AK1, HG002, and CHM13 (PacBio) datasets. Strikingly, those SNVs were likely to be shared among the four samples and even their frequencies were strongly correlated (Figure 3e). Similar patterns were confirmed for chromosomes 17 and X as well (Supplementary Figures 8, 9). One might expect that the pairs of Asian samples (B561-AK1) or European samples (HG002-CHM13) would show stronger correlations than the geographically unrelated pairs. However, we did not observe such a pattern, and AK1-CHM13 was always the most strongly correlated pair across the three chromosomes we analyzed (Supplementary Figure 10).

**Figure 3.**
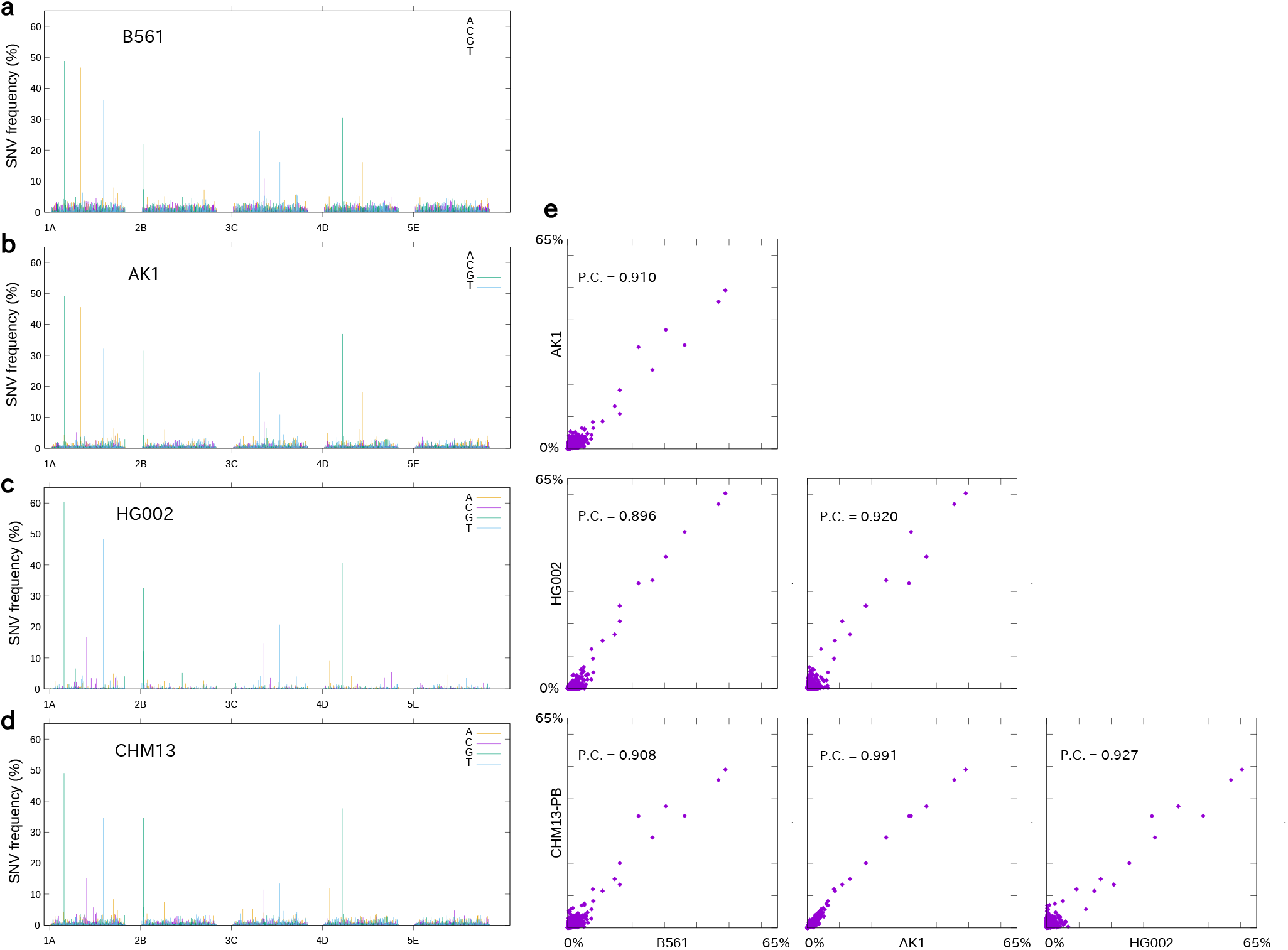
Single-nucleotide variants found in the 5-mer canonical HOR of chromosome 11 in four samples. (a-d) The frequency (y-axis) of alternative bases according to positions over the 5-mer canonical HOR (x-axis). Data for each alternative base is colored in yellow, purple, green, or blue if it is A, C, G, or T, respectively. (e) Correlation of frequency of variants among four samples. Each point represents a specific SNV, and plotted according to its frequency in two samples (x-axis and y-axis). P.C = the Pearson correlation between frequencies of all variants.

We found that the ONT read dataset for CHM13 was the most dissimilar to all the PacBio datasets (Supplementary Figures 10, 11, 12); thus, these reads might be unsuitable for SNV analysis, presumably due to persistent systematic sequencing errors.

Overall, we observed the conservation of SNVs in canonical HORs, in contrast to the diversity of HOR patterns.

## Discussion

In this study, we cataloged variant HORs found in the diploid human centromeres of three chromosomes 11, 17, and X by effectively utilizing long read data without explicit sequence assembly. We expanded the knowledge of variant HORs that have been documented^20–22^, and comparison of the variant HOR frequencies among four individuals revealed a substantial diversity in human centromeres. For some cases, we could even trace structural changes among samples via rare and unique motifs, *i.e.*, relatively rare combinations of variant HORs. By analysing three diploid samples, we detected several loci that appeared to represent alternative alleles, but none of them were the same, indicating that such variations are far from fixed among human populations and that other configurations remain to be discovered. Here, we note that regions harboring variant HORs were only a portion of the entire picture. Assuming that similar structural changes are occurred within the sea of tandem replicates of canonical HORs, there is even greater diversity in centromeres than we conservatively estimated here. Thus, we hypothesize that the tandem nature of centromeric arrays makes them extremely variable, as with other tandem repeats, and if one could accurately measure them, they would contain sufficient information to distinguish individual persons just as microsatellites can. We would be able to prove or disprove the hypothesis by exploring how diverse the structures of HOR arrays may be within a single population.

While centromeres were highly variable at the HOR level, we demonstrated that the frequent SNVs in the HOR units were clearly conserved among diverse populations, suggesting that they have spread before the geographical separation of the human population. These seemingly contradictory observations can be reconciled easily, because the unequal crossovers and gene conversions that would be responsible for the introduction of HOR-level structural variation can contribute to homogenization at the SNV level. What does it means to have such large structural diversity in centromeres? Because centromeres have a fundamental importance to proper chromosome segregation during cell division, it was once considered surprising to observe great diversity in centromeric sequences across different eukaryotic taxa (“centromere paradox”)^42^. Centromere drive theory explained the rapid evolvability of centromeres via genetic conflict during female meiosis I, rendering the centromeres as a crux of the molecular identity of species^43^. Nevertheless, growing evidence suggests that centromeres can be highly variable within a single species^5, 10, 20, 24^, and our findings add another layer of diversification, ultimately challenging the notion of stable human centromeres.

Several mechanisms can contribute to such structural diversity within centromeres: unequal crossover between sister chromatids, meiotic unequal crossover, gene conversion, and homologous recombination resulting in noncrossover products, to name a few. Among them, meiotic crossovers might arguably be excluded as a major driving force because they are suppressed near centromeric regions^7, 44^, and consequently, centromeres are reported to form large conserved linkage-disequilibrium blocks^10^. On the one hand, the structural diversity within centromeres can be best explained by frequent unequal crossovers between sister chromatids as well as gene conversions. On the other hand, centromere integrity in a human population might have been maintained through occasional gene conversions and infrequent meiotic crossovers, both of which can counteract the diversification processes by effectively homogenizing sequences among different alleles. Notably, all these mechanisms are consistent with the local, progressive expansion suggested in this study as well as in previous evolutionary analyses^45^. We speculate that all these mechanisms might have contributed to the current landscape of human centromeres.

Recently, in addition to a complete linear assembly of a human Y centromere, a telomere-to-telomere assembly spanning the whole human X centromere utilizing ultralong ONT reads has been reported, showing, at last, the time is ripe to investigate centromeres in terms of sequencing technology^36, 37^. With an increasing number of individual genomes from the same or closely related populations sequenced by long reads, one would be able to precisely observe the processes of diversification and homogenization that occur within human centromeres. Therefore, such a study should provide a basis to delineate the complex mechanisms involved and to understand the true nature of centromere evolution.

## Methods

### Preparation of long read sequencing data

For SMRTbell library preparation, B cell DNA (named B561 in the main text) was sheared using a Diagenode’s Megaruptor 2 with software setting 75 kb and purified using a 0.6 × volume ratio of AMPure beads (Pacific Biosciences, Menlo Park, CA, USA). SMRTbell libraries for sequencing were prepared using the “Procedure & Checklist-Preparing *>* 30 kb Libraries Using SMRTbell Express Template Preparation Kit” protocol. Briefly, the steps included (1) DNA repair, (2) blunt ligation with hairpin adapters with the SMRTbell Express Template Preparation Kit (Pacific Biosciences) (3) 15 kb size cutoff size selection using the BluePippin DNA Size Selection System by Sage Science, and (4) binding to polymerase using Sequel Binding Kit 2.1, later Sequel Binding Kit 3.0, (Pacific Biosciences). SMRTbell libraries were sequenced on Sequel SMRT Cells (Pacific Biosciences) using diffusion loading, 30 kb-insert size and 600-min movies. Other long read data, *i.e.*, from AK1^30^, CHM13^31^, HG002(Hi-Fi)^33^, and CHM13 (ONT ultra-long)^37^ were obtained via public repositories respectively.

### Filtering out non-centromeric reads

To enrich the centromeric reads *in silico*, we calculated the reference 6-mer frequency vector with the non-redundant catalog of 1194 monomers. We also calculated the “query” 6-mer frequency vector (normalized by length in bp) and its dot product with the reference for each long read. The dot products exhibited a bi-modal distribution which represents the mixture of centromeric and non-centromeric reads. Thus, only reads with the dot product greater than 200 were included in later analysis. We modified squeakr^46^ to perform these steps.

### Monomer encoding of long reads

For monomer encoding of long reads, the catalog of human alphoid monomers was adapted from: https://raw.githubusercontent.com/volkansevim/alpha-CENTAURI/nhmmscan/example/MigaKH.HigherOrderRptMon.fa Then, the distinct 1197 monomers were mapped by blastn (version 2.4.0+) to long reads with following parameters:

~~~
-max_target_seqs 1000000 -word_size 7 -qcov_hsp_perc 60 -outfmt "17 SQ"
~~~

Optimal assignment was calculated via dynamic programming procedure, maximizing the following quantity:

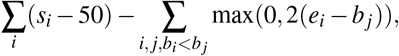

where *i* indexes monomers assigned to the read, *s*_*i*_ is the BLAST score of the hit, (*b*_*i*_, *e*_*i*_) the region covered by the monomer. Intuitively, it tries to assign as many monomers with acceptable scores as possible, due to the first term. The second term penalized the overlaps (*cf.* gaps were not penalized) so that each segment of the read be assigned at most one monomer.

### HOR encoding

Reads were then clustered based on the types of assigned monomers. First, each read was treated as a bag of monomers and represented as a vector encoding the relative frequencies of each type of monomer assigned to the read. Then, K-means clustering (*K* = 40) was performed with the cosine distance (*i.e.*, 1 − correlation) so that similar HOR patterns were represented in a single cluster. The three clusters of reads from chromosome 11, 17, and X can be identified, since they mainly comprised the characteristic (SF3, supra-chromosomal family 3) monomers. In each read cluster for chromosomes 11, 17, and X, recurrent combinations of monomers were identified as HORs. Within HOR patterns, no gap of *>*100 bp was allowed between neighboring monomers. With the list of identified HORs, reads were processed again to be encoded as series of assigned HORs plus the mismatches against the reference monomers.

**Table 1.** The probabilities that a given window contains multiple variant HORs, calculated for typical HOR unit lengths, variant HOR fractions, and window sizes. *de novo* assembly of centromeres solely based on variant HORs is possible if the centromeres are completely covered by long reads anchoring multiple variant HORs. The rightmost numbers in this table approximate the probability that a randomly sampled read meets such a criterion.

| HOR unit length (bp) | variant HOR fraction | Window size (bp) | Prob. >1 variants (%) |
| --- | --- | --- | --- |
| 855 = 171 * 5 (chr11) | 2% | 50,000 | 32.34 |
|  |  | 100,000 | 67.68 |
|  |  | 150,000 | 86.68 |
|  | 1% | 50,000 | 11.47 |
|  |  | 100,000 | 32.32 |
|  |  | 150,000 | 52.33 |
| 2,052 = 171 * 12 (chrX) | 2% | 50,000 | 8.26 |
|  |  | 100,000 | 24.94 |
|  |  | 150,000 | 43.03 |
|  | 1% | 50,000 | 2.39 |
|  |  | 100,000 | 8.34 |
|  |  | 150,000 | 16.58 |
| 2,736 = 171 * 16 (chr17) | 2% | 50,000 | 4.95 |
|  |  | 100,000 | 16.18 |
|  |  | 150,000 | 29.39 |
|  | 1% | 50,000 | 1.38 |
|  |  | 100,000 | 5.03 |
|  |  | 150,000 | 10.18 |

## Acknowledgements

The authors thank Drs. Wei Qu, Jun Yoshimura, Ms. Tsurutsuki Sugai, and Ms. Yoko Saito for sequencing B561 using PacBio Sequel. This study was supported in part by the Advanced Genome Research and Bioinformatics Study to Facilitate Medical Innovation from Japan Agency for Medical Research and Development (AMED) to S.M.

## Author contributions statement

Y.S. conceived and conducted the study. Y.S., G.M., and S.M. analysed the results and wrote the manuscript All authors reviewed the manuscript.

## Additional information

All sequencing data for B561 are available under the accession code JGAS00000000173 in the DDBJ Japanese Genotype-phenotype Archive for genetic and phenotypic human data.

The snapshot of all custom codes used for the study is available here: https://github.com/hacone/hc_temporary/releases/tag/submission-v.0.9

The authors declare no competing interests.

---

**Supplementary Figure 1.**
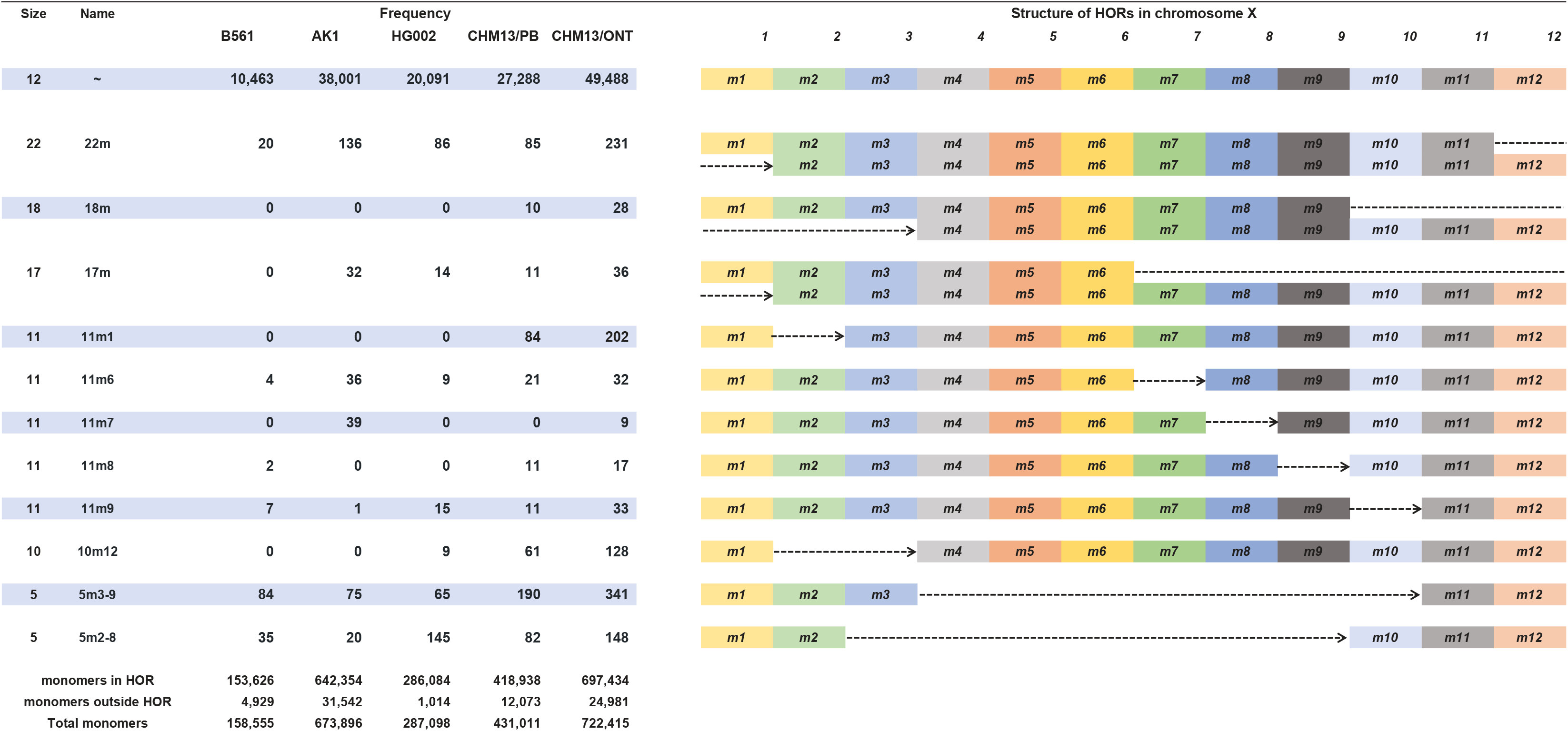
Detected HORs in chrX. The table summarizes the absolute frequencies of each type of HOR, detected in long reads of each sample. The structure of HORs are presented in the right hand side of each row. The rectangles represent the presence of alphoid monomers, whose name (e.g m1) are written on the rectangles. For example, variant HOR named "1Om12" is characterized by deletion of two monomers, m2 and m3, resulting in the 1O-mer (1.7kbp) HOR (no gap is allowed between m1 and m4 to be detected as 1Om12 variant. The numbers of alphoid monomers which were detected in all reads, which were organized into HORs (including truncated canonical HORs found at the edges of reads), and which were not, are presented in the last row of the left table.

**Supplementary Figure 2.**
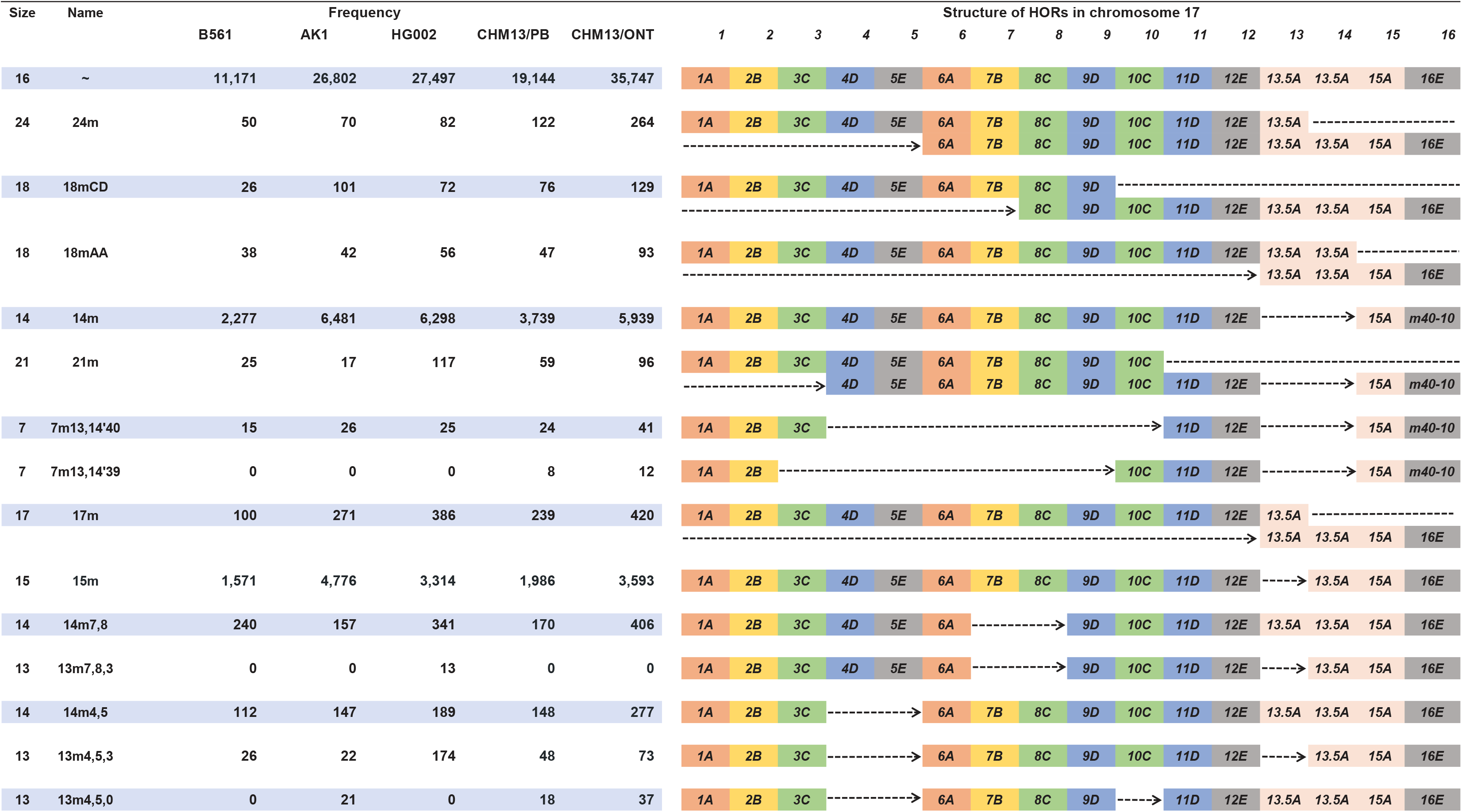
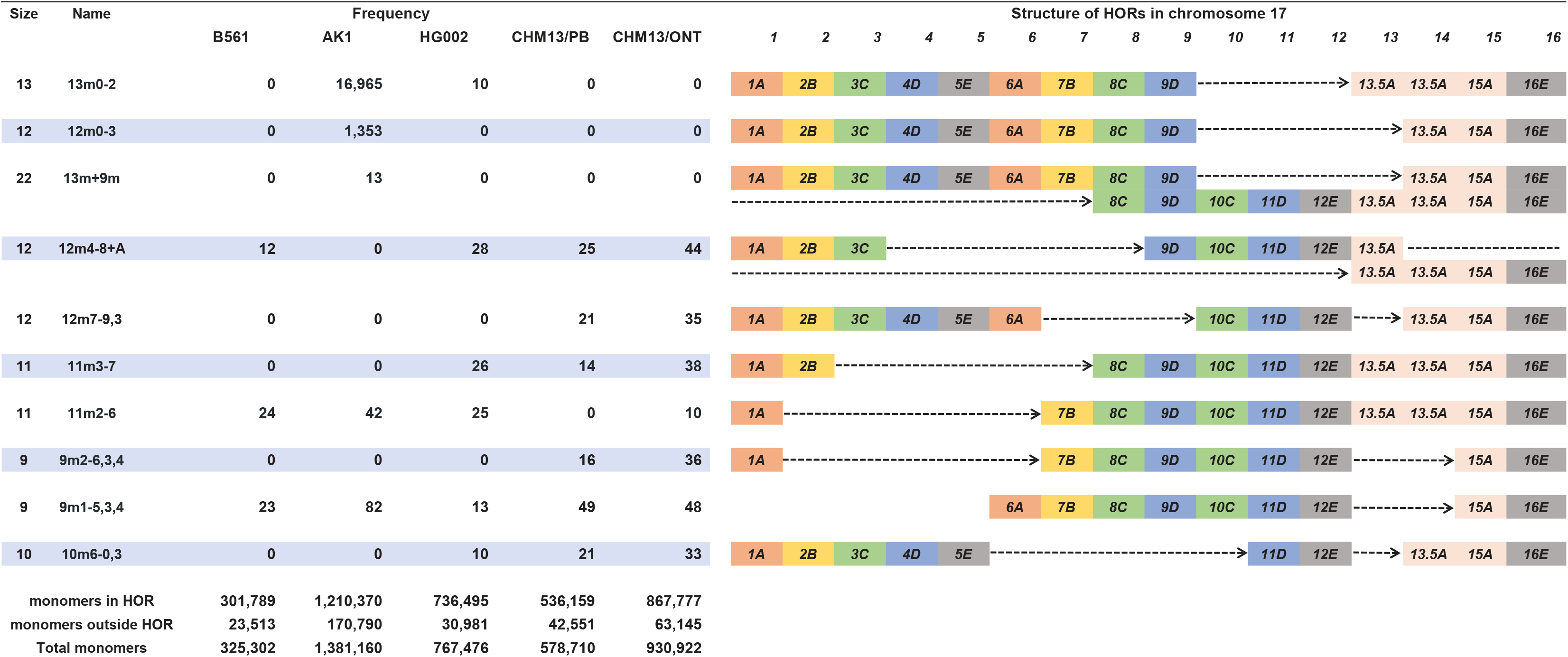
Detected HORs in chr17. The table summarizes the absolute frequencies of each type of HOR, detected in long reads of each sample. The structure of HORs are presented in the right hand side of each row. The rectangles represent the presence of alphoid monomers, whose names (e.g 1A) are written on the rectangles as in Supplementary Figure 1. Rectangles (monomers) are colored according to subcategory to which they belong (see Alexandrov et al. 2001). In HOR detection, the 13th and 14th monomers were identified, for they were too similar to be distinguishable in row long reads.

**Supplementary Figure 3.**
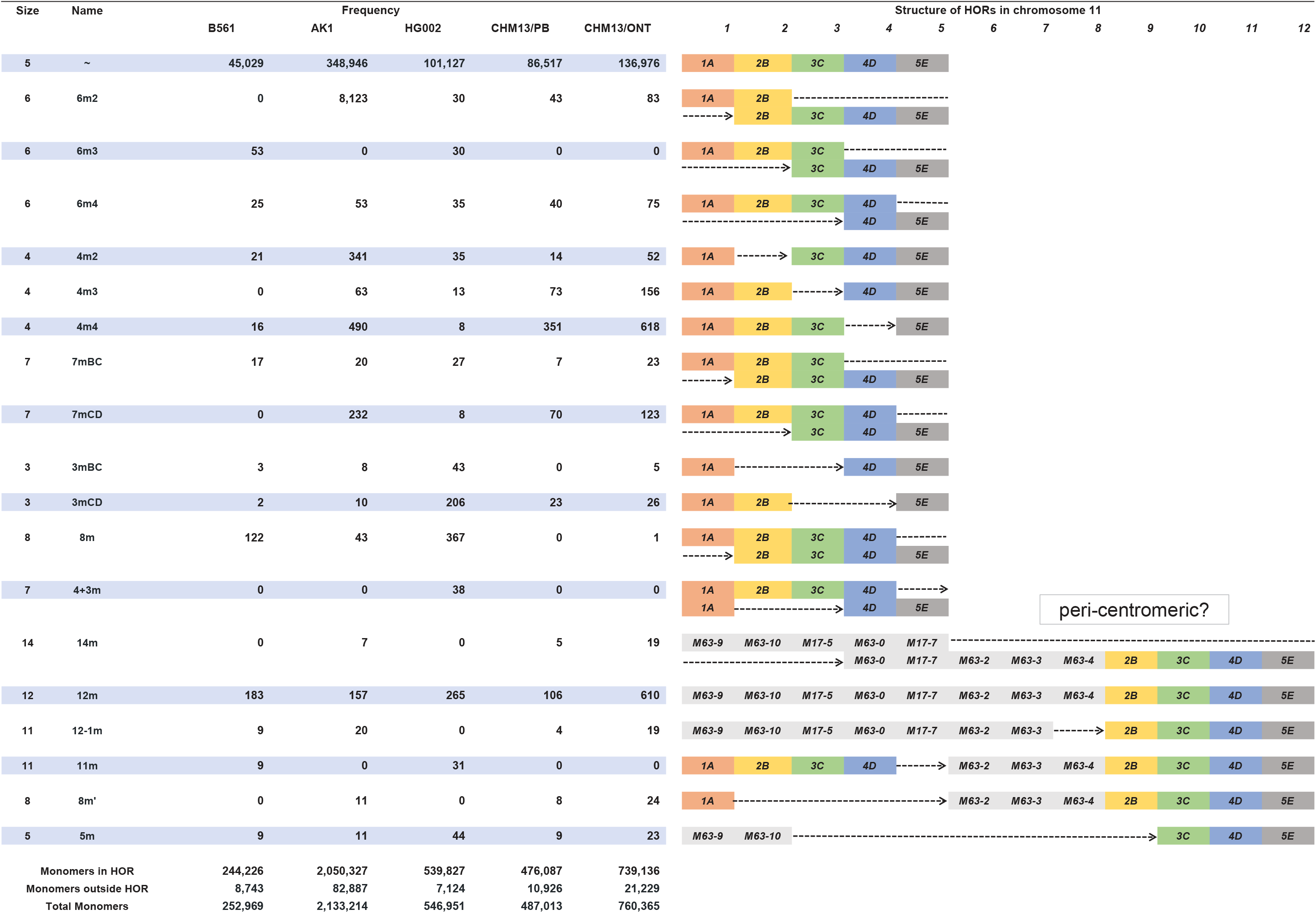
Detected HORs in chr11. The table summarizes the absolute frequencies of each type of HOR, detected in long reads of each sample. The structure of HORs are presented in the right hand side of each row. The rectangles represent the presence of alphoid monomers, whose names (e.g 1A) are written on the rectangles, and rectangles (monomers) are colored according to subcategory to which they belong (see Al exandrov et al. 2001) as in Supplementary Figure 2.

**Supplementary Figure 4:**
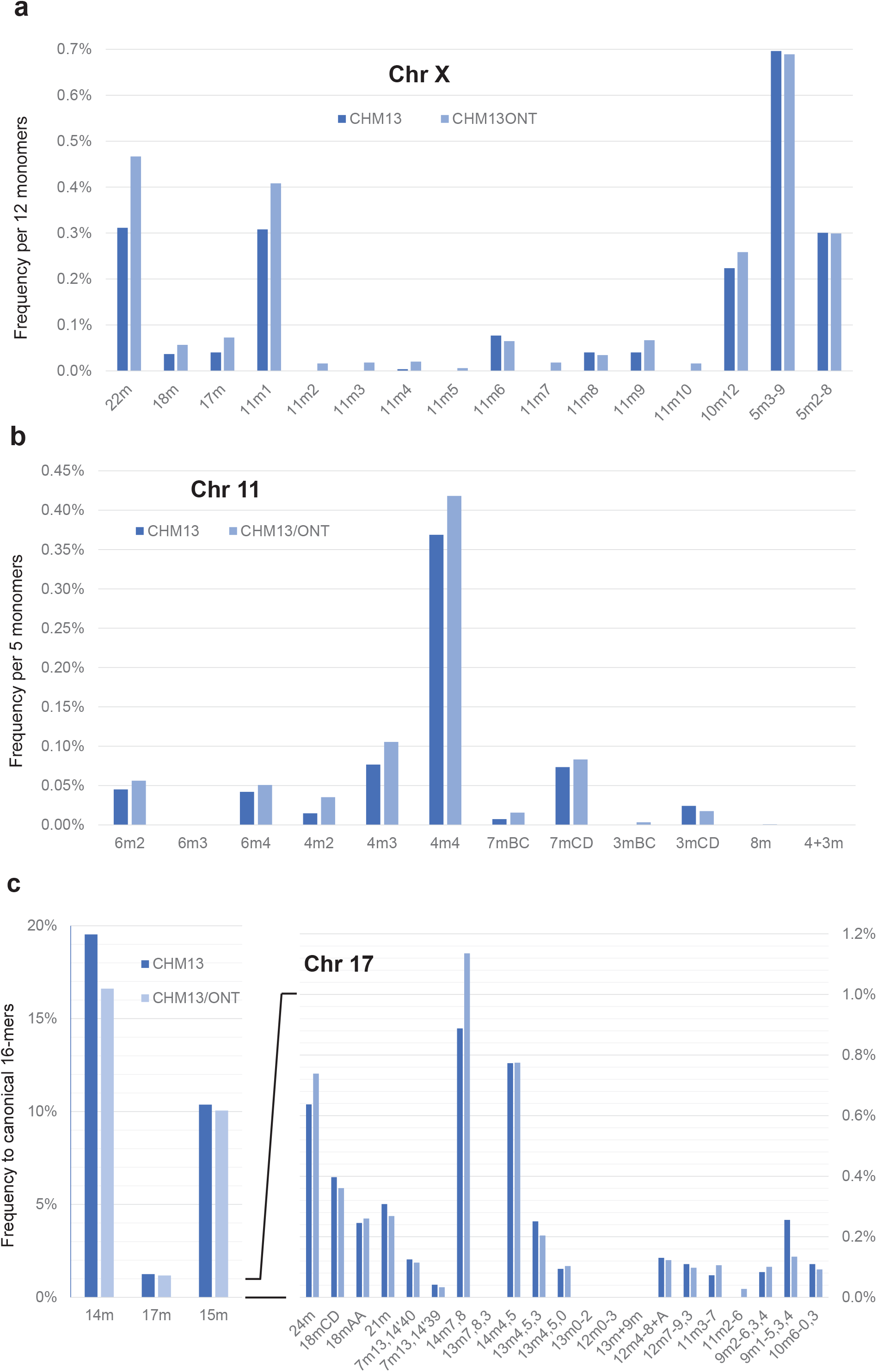
Relative frequencies of detected HORs for CHM13. The plot shows the comparison between PacBio reads (CHM13: darker blue) and ONT reads (CHM13ONT: lighter blue), for (a) chromosome 11, (b) chromosome 17, and (c) chromosome X. The definitions of canonical or variant HORs are the same as in Supplementary Figures 1–3.

**Supplementary Figure 5.**
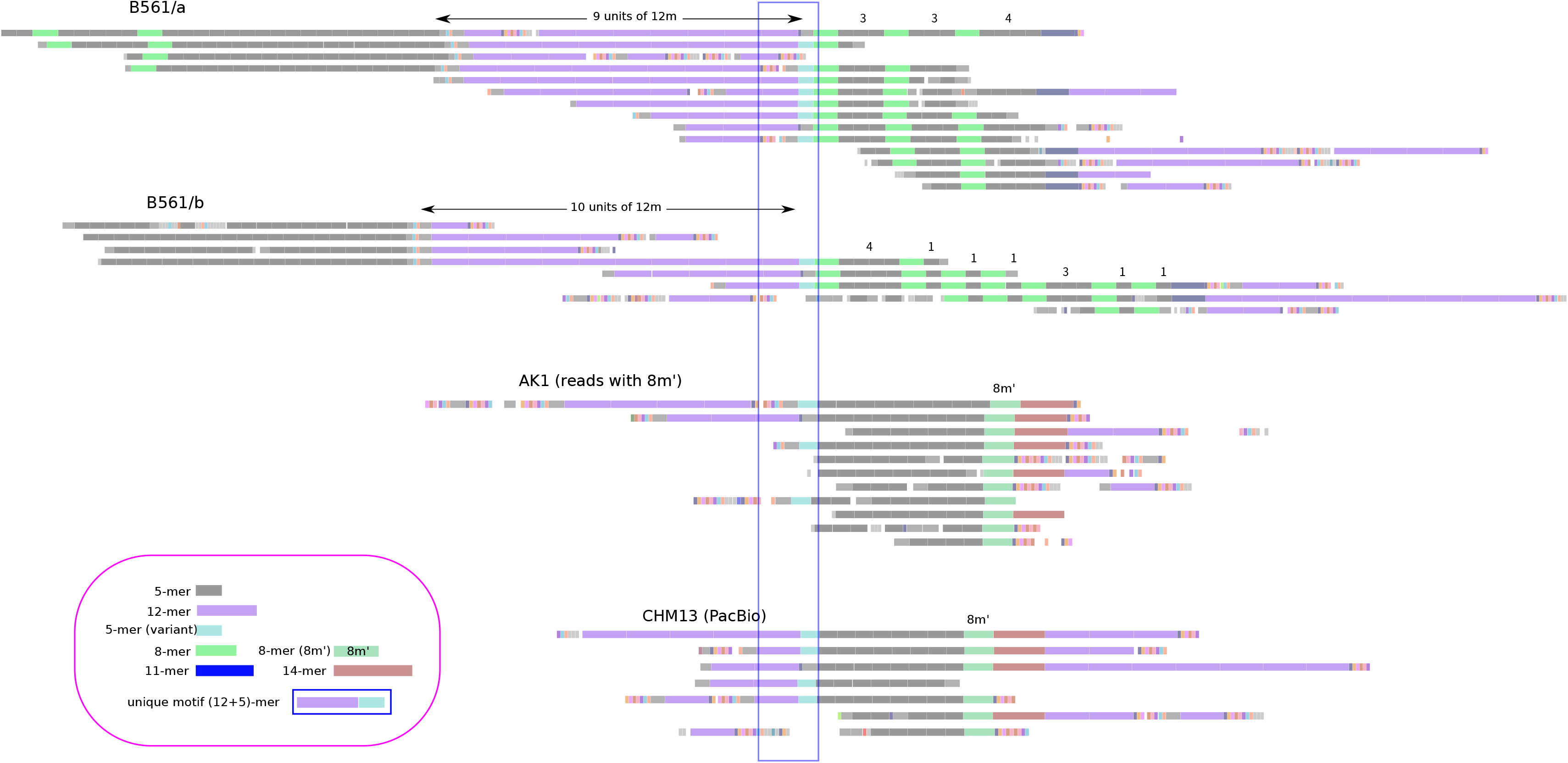
Visualization of HOR-encoded long reads with a motif. Each line represents a single HOR-encoded long read. The 5-mer canonical HOR for chromosome 11 and the monomers in the canonical HOR are represented by grey rectangles. The other variant HORs and monomers are colored differently. HORs and monomers are drawn at the positions proportional to their actual coordinates in long reads. The blue line at the center specified the motif position (a combination of 12-mer and 5-mer variants). B561 and HG002 (Supplementary Figure 6) shared similar downstream patterns while AK1 and CHM13 share the downstream patterns with 8-mer variant (8m’, with the prime). All reads with the motif are shown here.

**Supplementary Figure 6.**
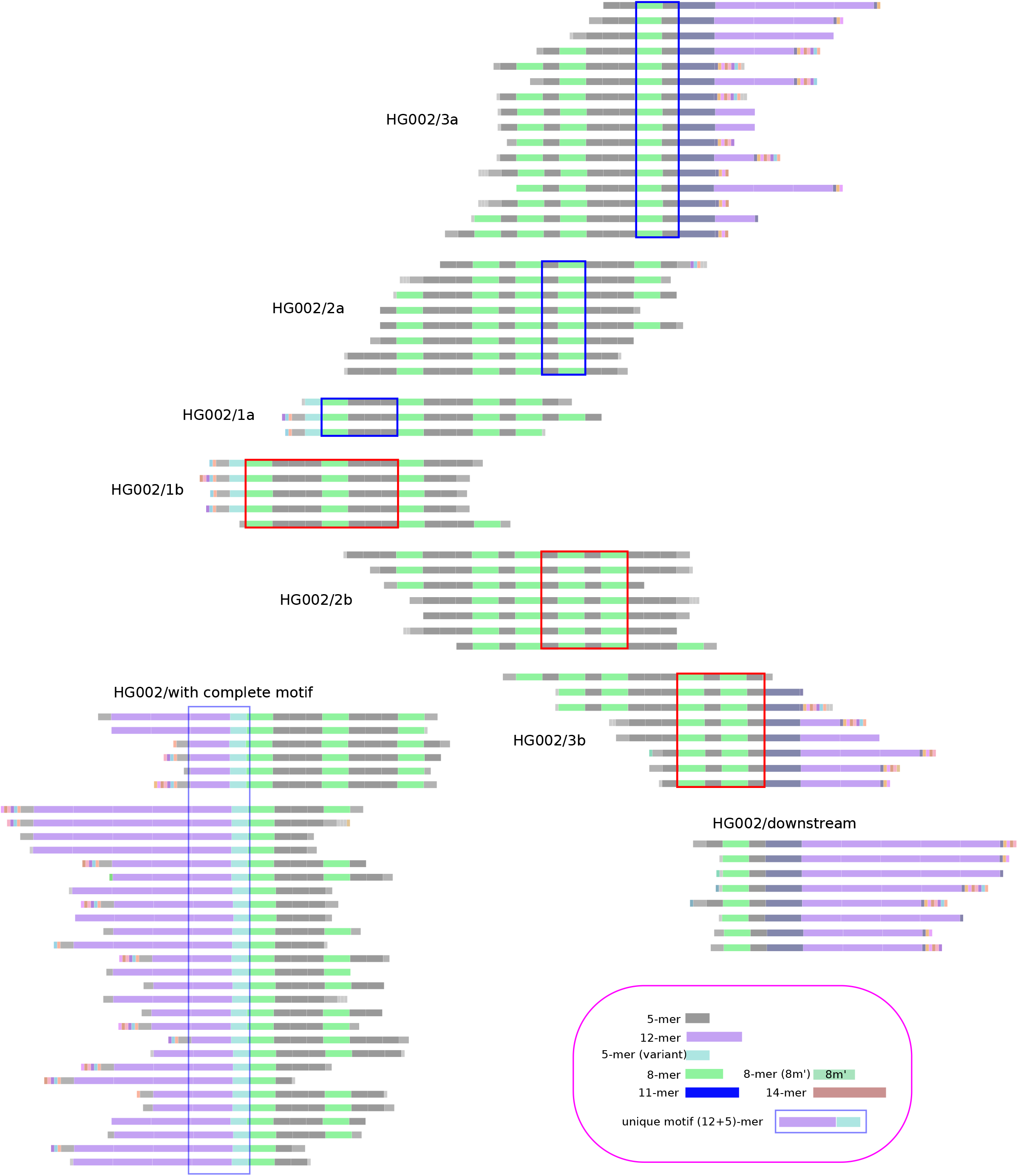
Visualization of HOR-encoded long reads with a motif in HG002. Each line represents a single HOR-encoded long read. The 5-mer canonical HOR for chromosome 11 and the monomers in the canonical HOR are represented by grey rectangles. The other variant HORs and monomers are colored differently. HORs and monomers are drawn at the positions proportional to their actual coordinates in long reads. The blue line in the left cluster specifies the motif position (a combination of 12-mer and 5-mer variants). As with B561 (Supplementary Figure 5), HG002 is characterised by the patterns with 8-mer variants, but the downstream patterns appeared complex and at least two alternative structures were suggested. All reads with the motif are shown here.

**Supplementary Figure 7.**
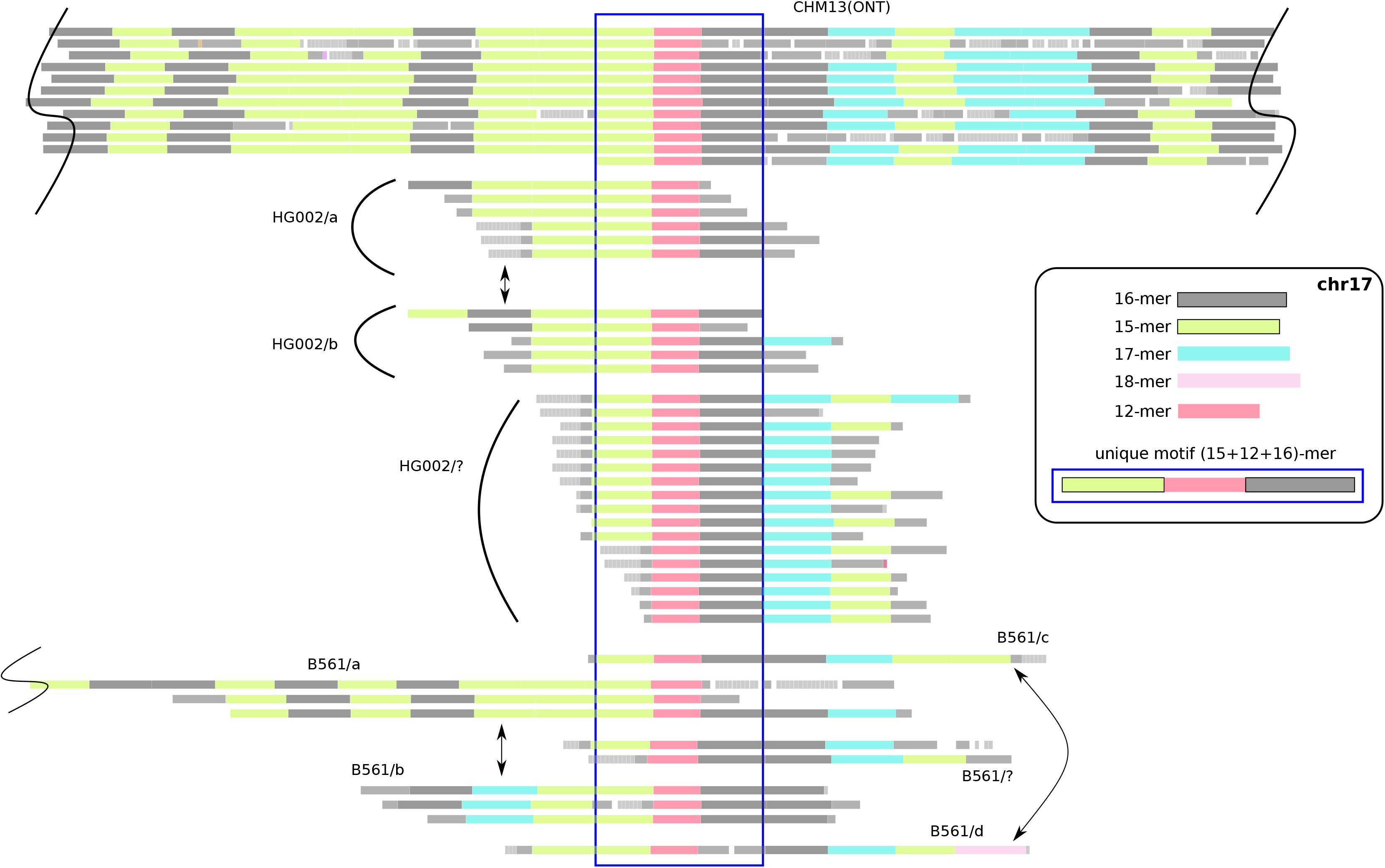
Visualization of HOR-encoded long reads with a motif. Each line represents a single HOR-encoded long read. Only a portion of the read is shown when the whole read does not fit in the figure. The 16-mer canonical HOR for chromosome 17 and the monomers in the canonical HOR are represented by grey rectangles. The other variant HORs and monomers are colored differently. HORs and monomers are drawn at the positions proportional to their actual coordinates in long reads. The blue line in the left cluster specifies the motif position (a combination of 15-mer variant, 12-mer variant, and canonical 16-mer). For B561 and HG002, all reads with the motif are shown here.

**Supplementary Figure 8.**
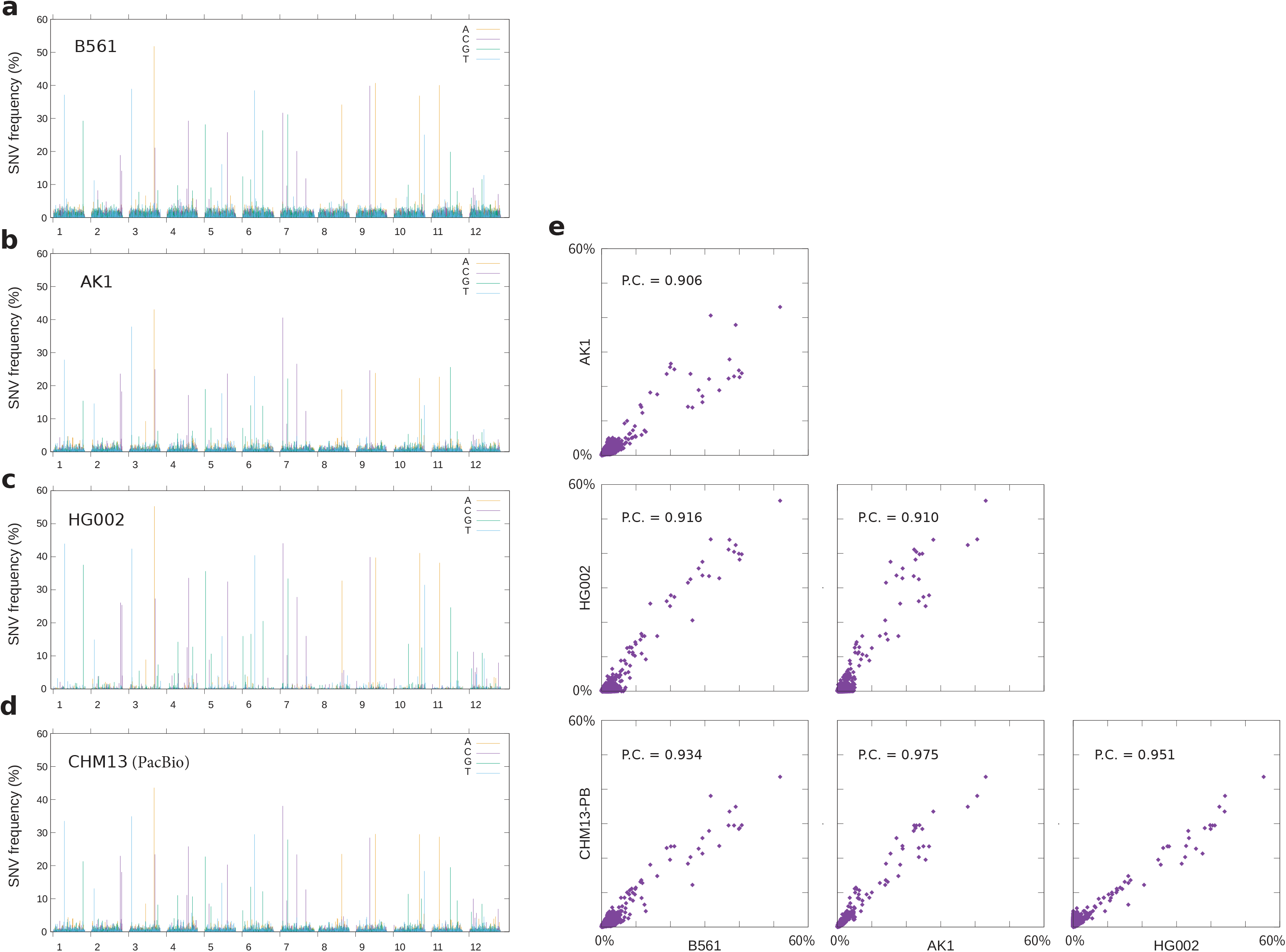
Single-nucleotide variants found in the 12-mer canonical HOR of chromosome X in four samples. (a-d) The frequency (y-axis) of alternative bases according to positions over the 12-mer canonical HOR (x-axis). Data for each alternative base is colored in yellow, purple, green, or blue if it is A, C, G, or T, respectively. (e) Correlation of frequency of variants among four samples. Each point represents a specific SNV, and plotted according to its frequency in two samples (x-axis and y-axis). P.C = the Pearson correlation between frequencies of all variants.

**Supplementary Figure 9.**
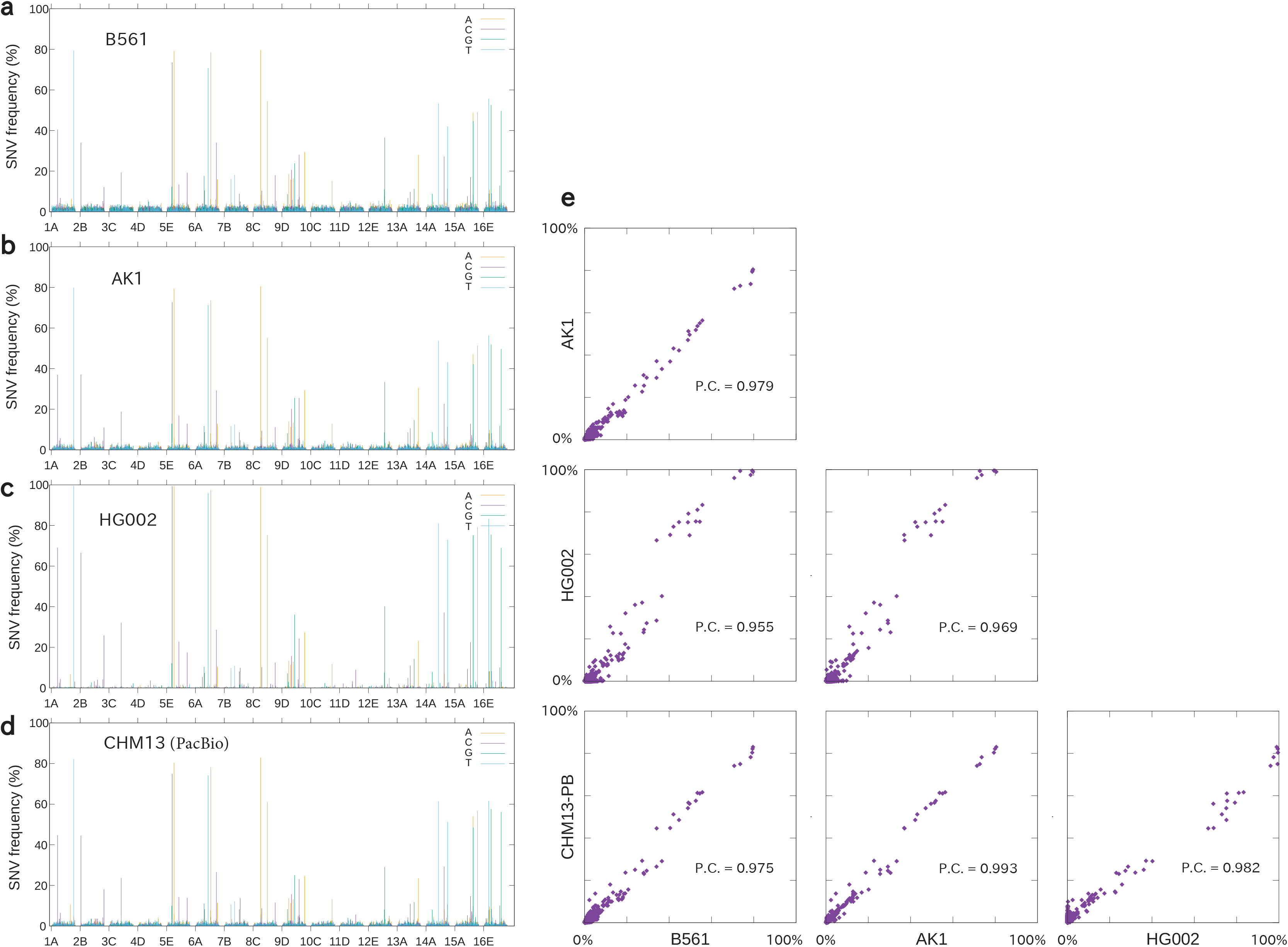
Single-nucleotide variants found in the 16-mer canonical HOR of chromosome 17 in four samples. (a-d) The frequency (y-axis) of alternative bases according to positions over the 16-mer canonical HOR (x-axis). Data for each alternative base is colored in yellow, purple, green, or blue if it is A, C, G, or T, respectively. (e) Correlation of frequency of variants among four samples. Each point represents a specific SNV, and plotted according to its frequency in two samples (x-axis and y-axis). P.C = the Pearson correlation between frequencies of all variants.

**Supplementary Figure 10.**
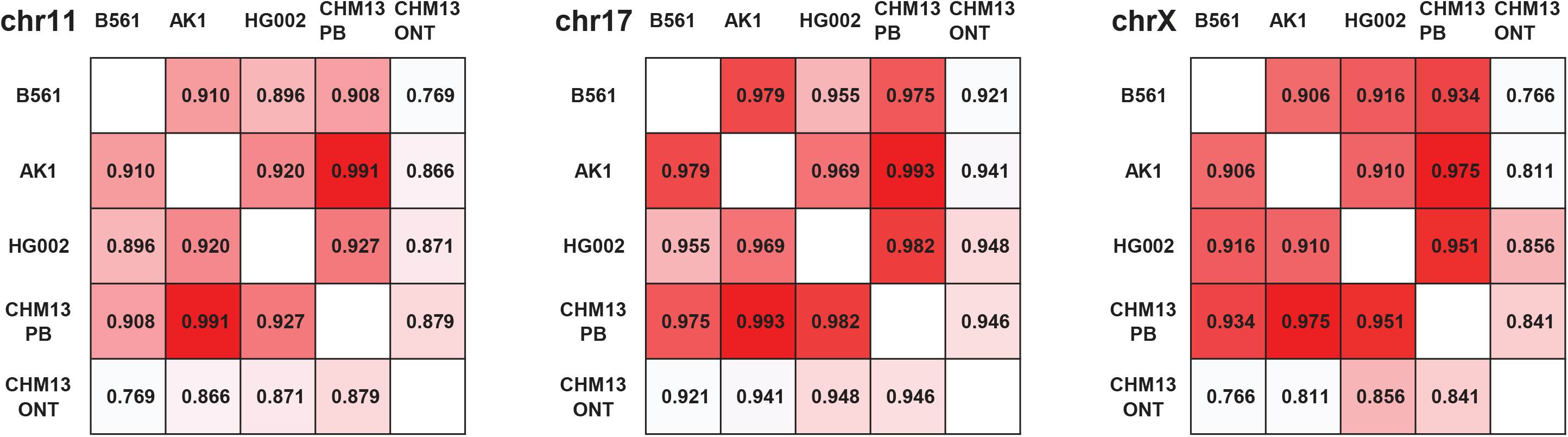
Pearson correlation coefficients for SNV frequencies. Data for three chromosomes, 11, 17, and X, for five datasets, B561, AK1, HG002, CHM13(PacBio), and CHM13(ONT) are shown. All variants were used to calculate Pearson correlation. The color gradation patterns were selected differently for each panel to show the least similar pairs to be white, and the most similar pairs to be solid red (the diagonals are left blank).

**Supplementary Figure 11.**
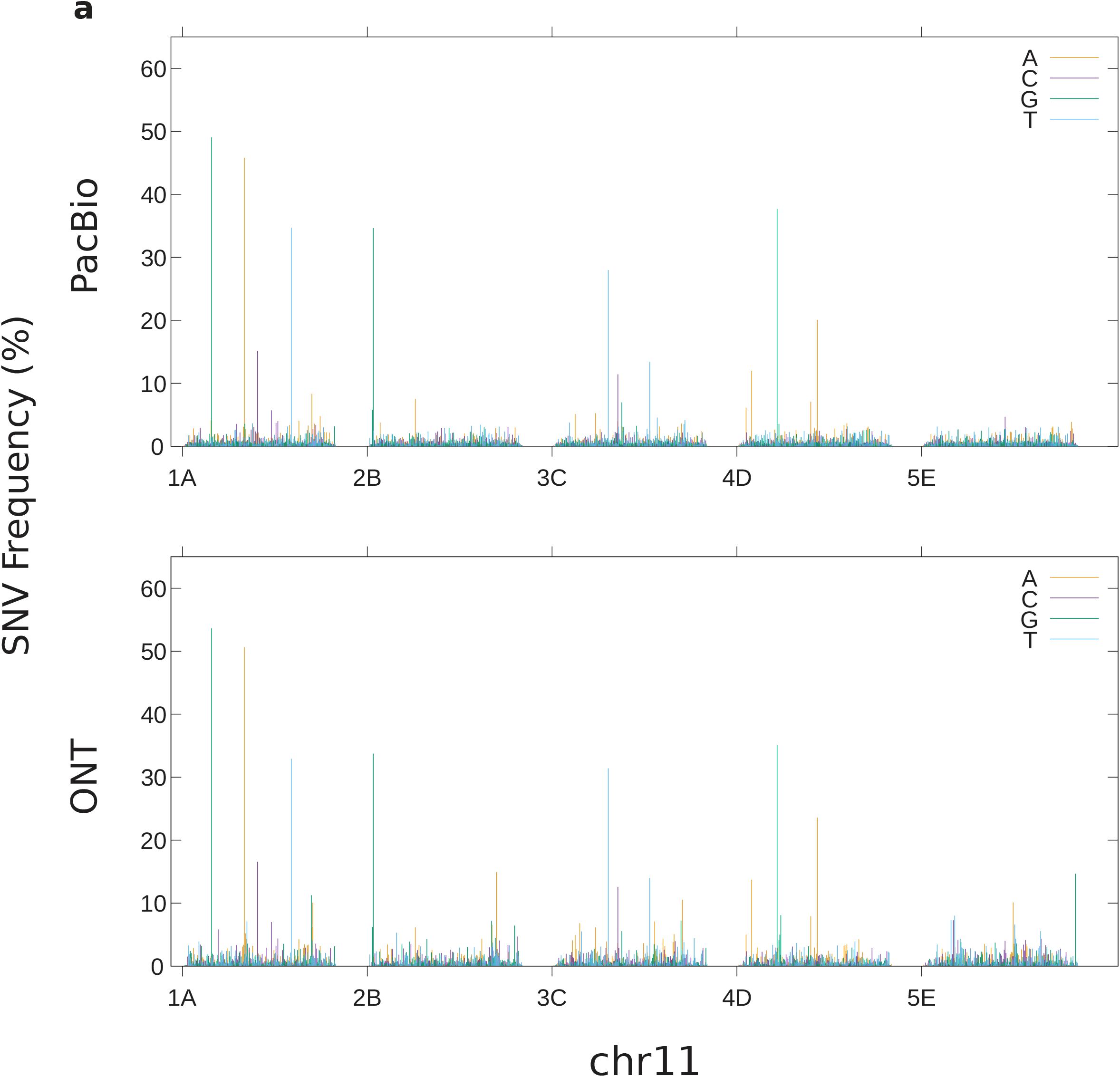
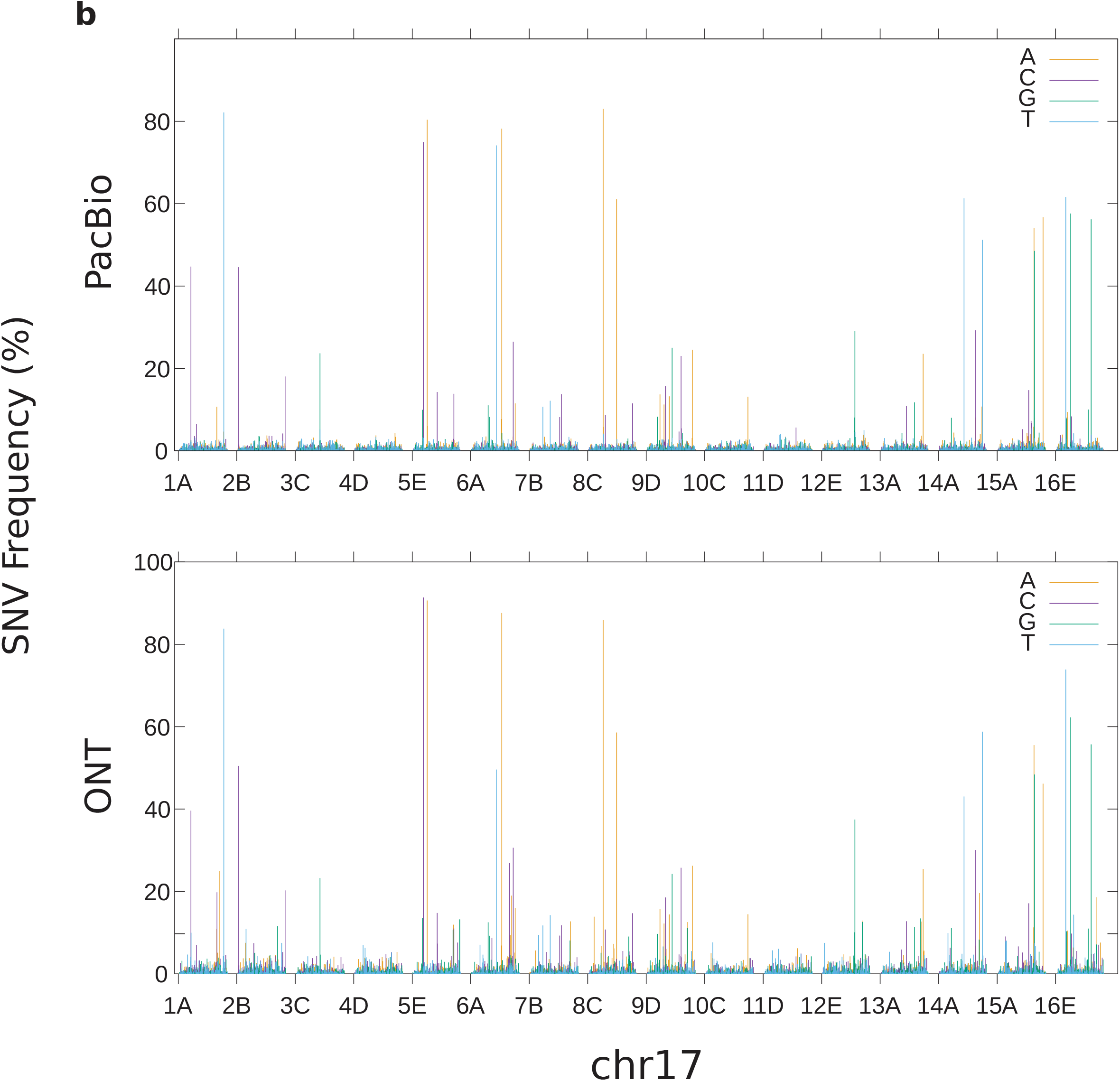
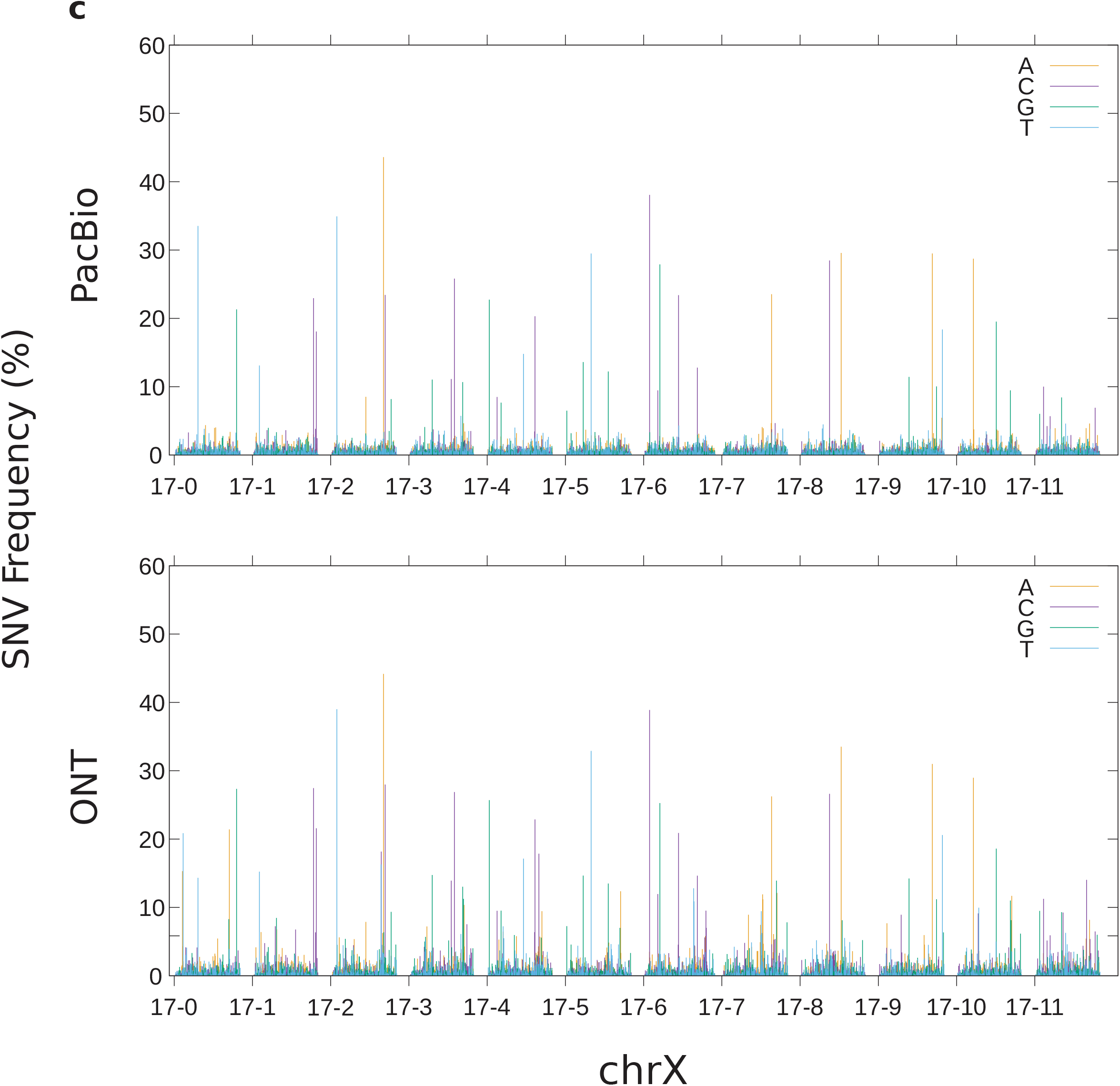
Single-nucleotide variants found in the canonical HORs of (a) chromosomes 11, (b) 17, and (c) X in CHM13, detected with PacBio or ONT. The frequency (y-axis) of alternative bases according to positions over the canonical HOR (x-axis). Data for each alternative base is colored in yellow, purple, green, or blue if it is A, C, G, or T, respectively.

**Supplementary Figure 12.**
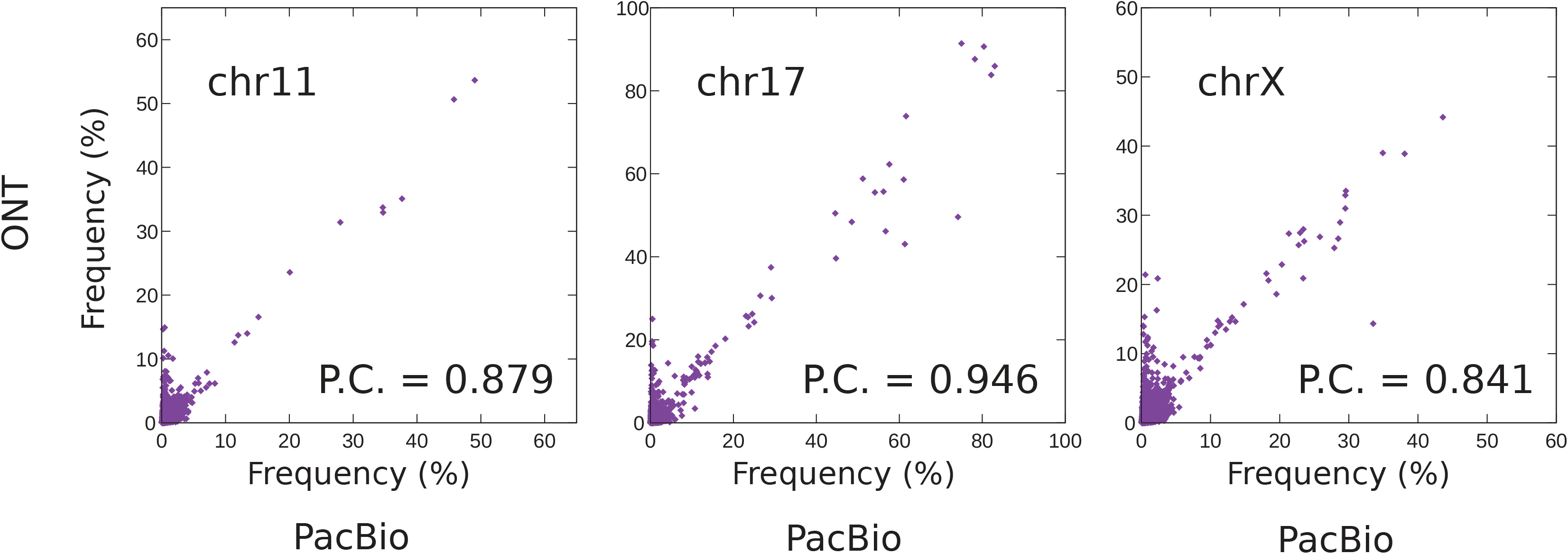
Correlation of frequency of detected variants between two sequencing technologies. Each point represents a specific SNV, and plotted according to its frequency in each dataset (x-axis for PacBio and y-axis for ONT). Data for three canonical HORs, 5-mer in chromosome 11, 16-mer in chromosome 17, and 12-mer in chromosome X, is shown. There are some spurious variants called only in ONT along the y-axis. P.C = the Pearson correlation between frequencies of all variants.

**Supplementary Table 1.** Summary of chi-square tests for distribution of variant HORs across four samples, in each of the three chromosomes X, 11, and 17. All statistical tests were performed based on the numbers shown in the tables in Supplementary Figure 1–3. For each type of variant HOR, we performed five tests in total: one for the observed number in each sample (degree of freedom = 1) and one for distribution across the four samples (degree of freedom = 4). The cells are colored in yellow if the minus log p-value exceeds the threshold for family-wise error rate of 0.01, which is shown below each table. The cells are colored in gray if p-value was too small to calculate its logarithm.

| chrX | minus log p-value |  |  |  |  |
| --- | --- | --- | --- | --- | --- |
|  | B561 | AK1 | HG002 | CHM13 | (all) |
| 22m | 2.07 | 0.24 | 1.46 | 0.40 | 1.83 |
| 18m | 0.53 | 1.33 | 0.83 | 4.65 | 4.32 |
| 17m | 1.90 | 1.32 | 0.26 | 0.71 | 1.79 |
| 11m1 | 2.61 | 8.10 | 4.57 | 33.97 | 43.80 |
| 11m2 | 0.34 | 1.49 | 0.51 | 0.63 | 0.97 |
| 11m3 | 0.19 | 0.76 | 0.29 | 0.35 | 0.26 |
| 11m4 | 0.13 | 0.28 | 0.19 | 0.74 | 0.19 |
| 11m5 | 0.13 | 0.47 | 0.19 | 0.23 | 0.08 |
| 11m6 | 0.73 | 0.93 | 0.86 | 0.09 | 0.77 |
| 11m7 | 1.41 | 8.67 | 2.37 | 3.06 | 11.40 |
| 11m8 | 0.20 | 1.63 | 1.01 | 3.83 | 3.80 |
| 11m9 | 1.06 | 3.17 | 2.50 | 0.17 | 3.97 |
| 11m10 | 0.00 | 0.00 | 0.00 | 0.00 | 0.00 |
| 10m12 | 2.24 | 6.86 | 0.86 | 19.44 | 24.75 |
| 5m3-9 | 8.11 | 11.46 | 1.71 | 10.51 | 26.70 |
| 5m2-8 | 0.35 | 17.41 | 28.25 | 0.07 | 41.59 |

minus log 0.01 / 16 = 3.20411998 minus log 0.01 / 64 = 3.80617997
| chr11 | minus log p-value |  |  |  |  |
| --- | --- | --- | --- | --- | --- |
|  | B561 | AK1 | HG002 | CHM13 | (all) |
| 6m2 | 139.29 |  | 298.22 | 248.02 |  |
| 6m3 | 74.66 | 11.77 | 4.38 | 3.35 | 88.12 |
| 6m4 | 3.88 | 4.29 | 0.99 | 3.52 | 8.76 |
| 4m2 | 1.26 | 8.74 | 4.79 | 8.78 | 18.90 |
| 4m3 | 3.17 | 2.28 | 1.95 | 26.45 | 29.05 |
| 4m4 | 9.33 | 0.69 | 30.44 | 84.81 | 119.02 |
| 7mBC | 6.03 | 3.27 | 4.52 | 0.56 | 10.42 |
| 7mCD | 6.02 | 3.13 | 9.39 | 3.36 | 17.21 |
| 3mBC | 0.25 | 4.74 | 27.27 | 2.34 | 30.06 |
| 3mCD | 3.94 | 28.35 | 141.05 | 1.50 | 168.38 |
| 8m | 35.63 | 53.18 | 178.44 | 18.24 | 277.58 |
| 4+3m | 1.06 | 5.74 | 33.58 | 1.76 | 37.25 |

minus log 0.01 / 12 = 3.07918125 minus log 0.01 / 48 = 3.68124124
| chr17 | minus log p-value |  |  |  |  |
| --- | --- | --- | --- | --- | --- |
|  | B561 | AK1 | HG002 | CHM13 | (all) |
| 24m | 0.57 | 2.89 | 1.63 | 7.89 | 9.25 |
| 18mCD | 1.06 | 0.86 | 1.18 | 1.09 | 1.68 |
| 18mAA | 2.31 | 1.44 | 0.19 | 0.42 | 2.00 |
| 14m | 4.39 | 11.09 | 1.83 | 14.46 | 26.73 |
| 21m | 0.32 | 9.43 | 7.38 | 0.77 | 14.00 |
| 7m13,14'40 | 0.44 | 0.19 | 0.36 | 0.38 | 0.17 |
| 7m13,14'39 | 0.52 | 0.95 | 0.97 | 5.38 | 4.77 |
| 17m | 2.22 | 1.91 | 3.27 | 0.44 | 4.63 |
| 15m | 0.40 | 70.87 | 13.71 | 35.93 | 114.69 |
| 14m7,8 | 27.28 | 13.87 | 2.12 | 1.87 | 39.91 |
| 13m7,8,3 | 0.72 | 1.37 | 4.71 | 1.06 | 4.70 |
| 14m4,5 | 3.76 | 2.63 | 0.13 | 0.59 | 4.25 |
| 13m4,5,3 | 0.97 | 11.19 | 19.48 | 1.03 | 27.99 |
| 13m4,5,0 | 1.63 | 1.86 | 3.43 | 2.70 | 6.01 |
| 13m0-2 |  |  |  |  |  |
| 12m0-3 | 40.01 |  | 96.90 | 67.81 |  |
| 13m+9m | 0.72 | 4.92 | 1.40 | 1.06 | 4.91 |
| 12m4-8+A | 0.61 | 5.24 | 0.87 | 2.14 | 5.59 |
| 12m7-9,3 | 1.02 | 2.00 | 2.05 | 13.04 | 14.03 |
| 11m3-7 | 1.67 | 3.43 | 3.51 | 1.00 | 6.06 |
| 11m2-6 | 3.26 | 1.85 | 0.40 | 5.24 | 7.21 |
| 9m2-6,3,4 | 0.84 | 1.61 | 1.65 | 10.11 | 10.43 |
| 9m1-5,3,4 | 0.08 | 4.20 | 7.67 | 1.17 | 9.60 |
| 10m6-0,3 | 1.37 | 2.76 | 0.01 | 6.89 | 7.74 |

minus log 0.01 / 25 = 3.39794001
minus log 0.01 / 100 = 4

